## Supplementary figures and images for "*Clostridioides difficile* exploits toxin-mediated inflammation to alter the host nutritional landscape and exclude competitors from the gut microbiota"

### Supplemental Figures

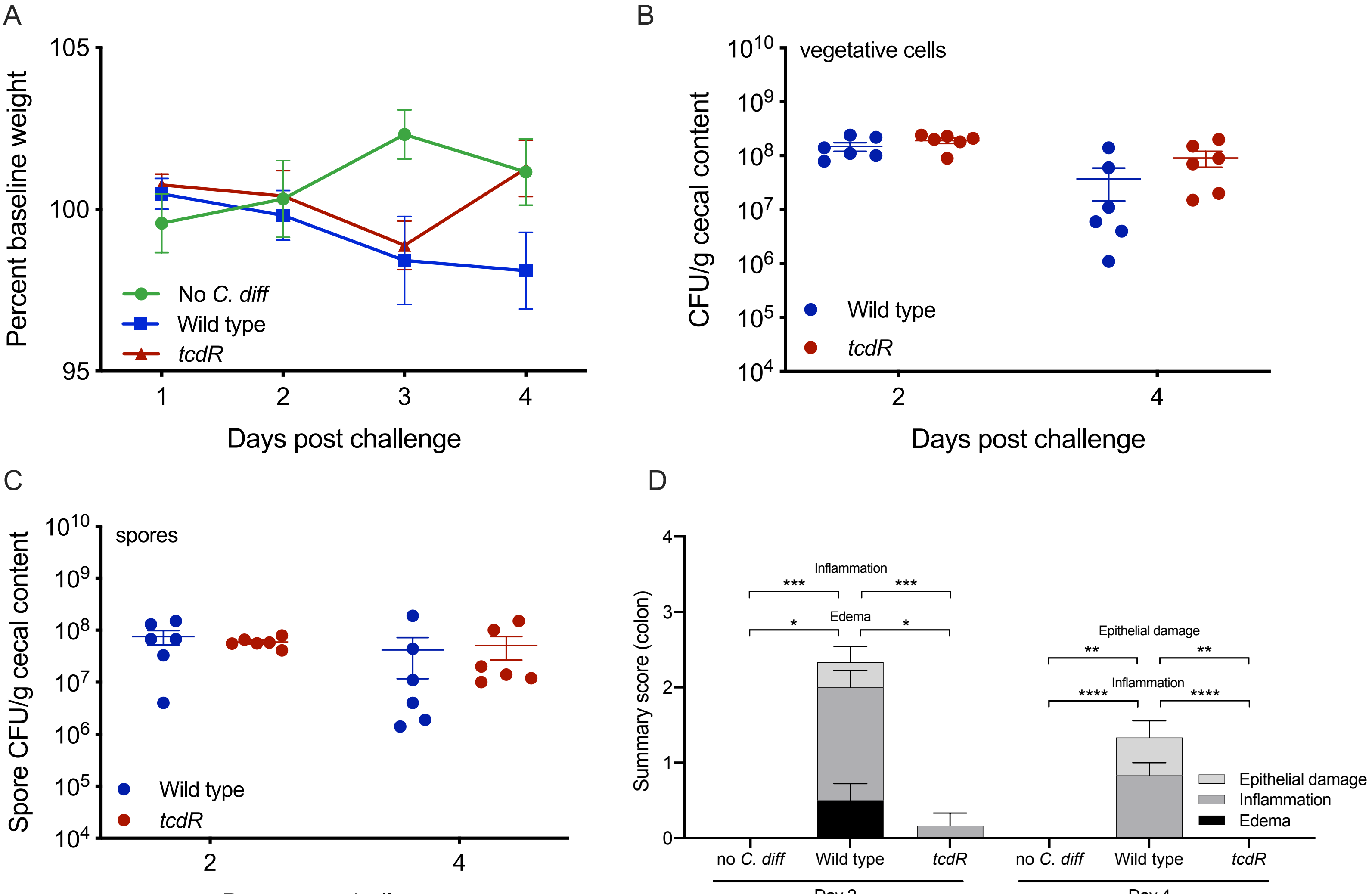

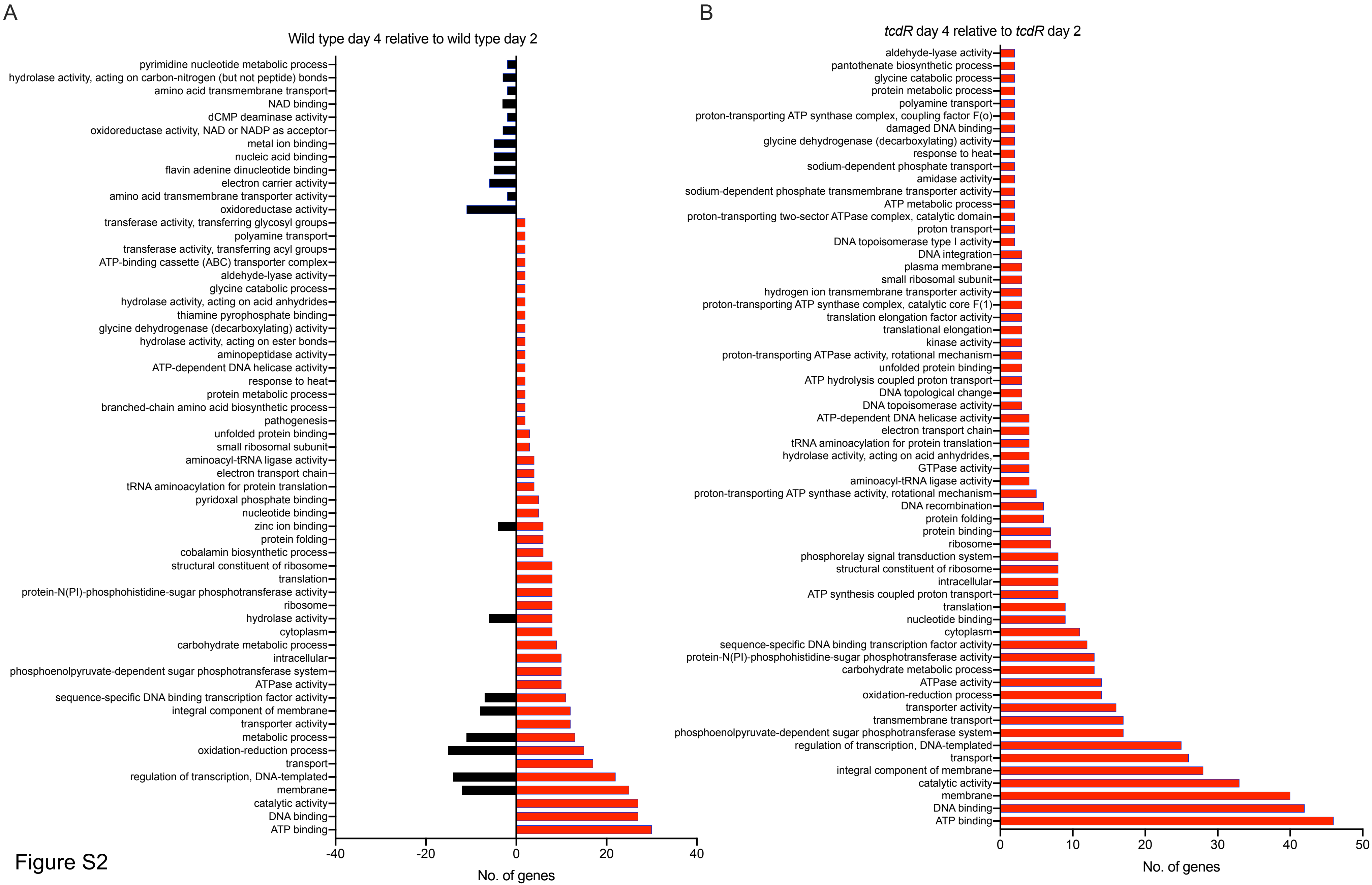

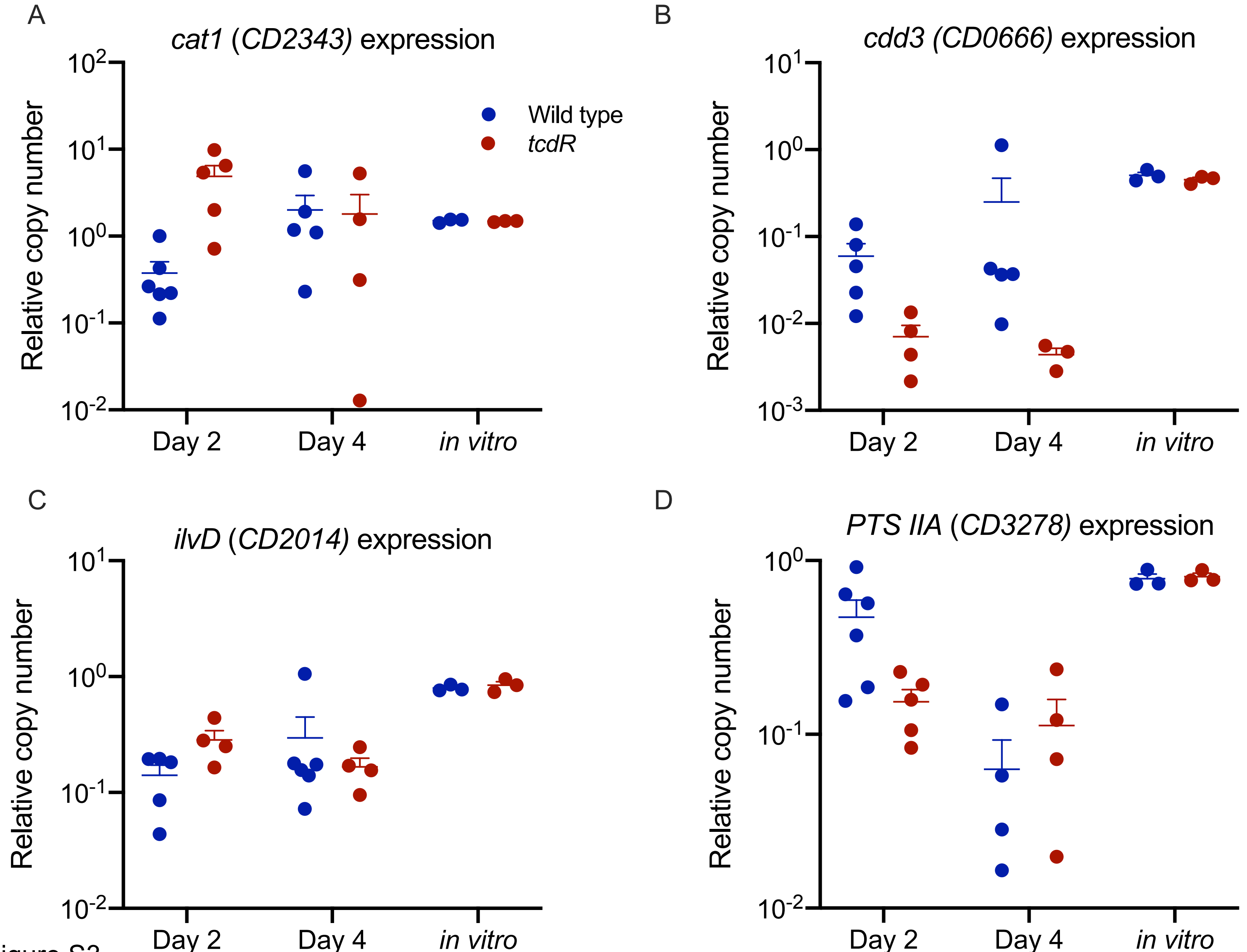

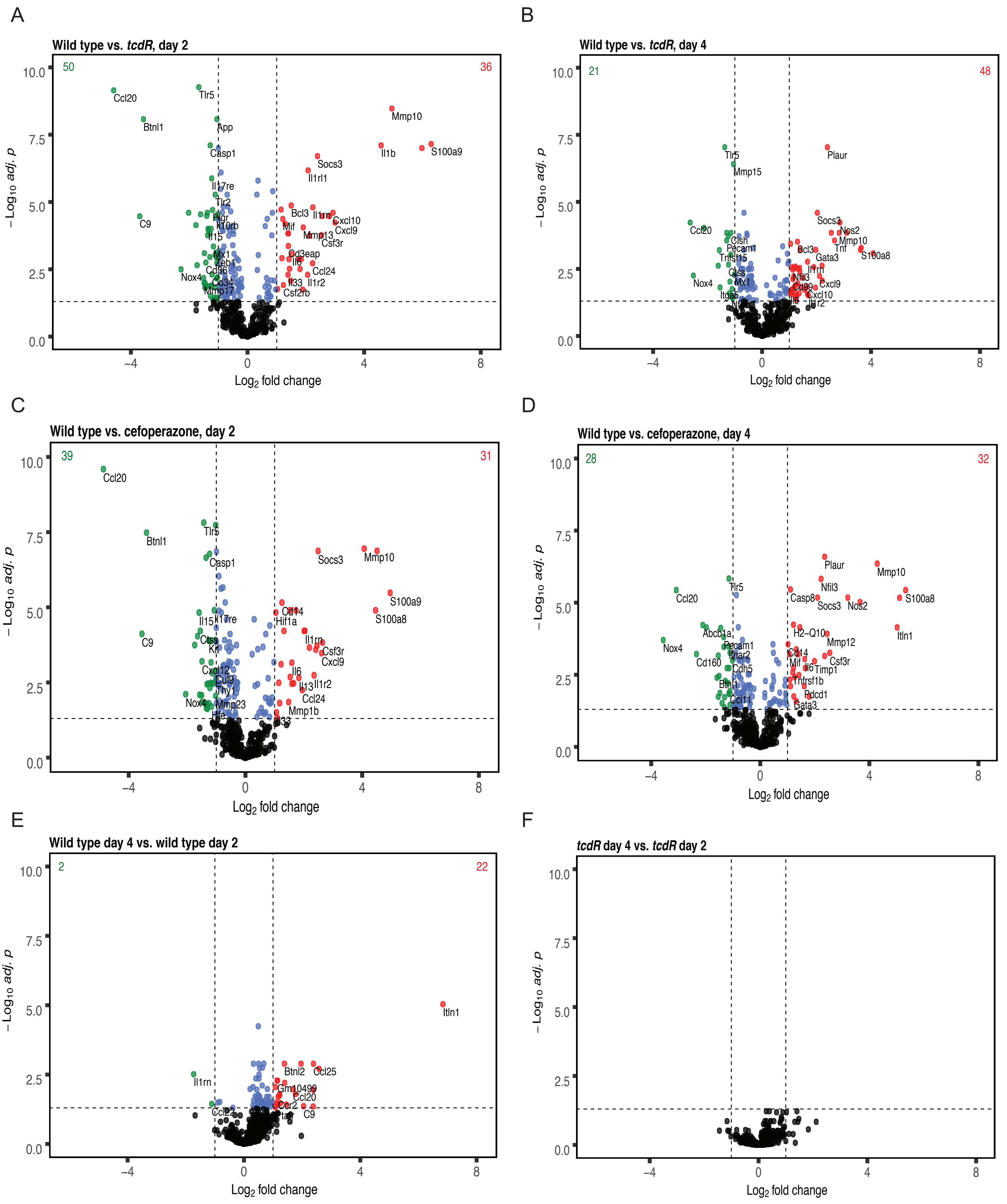

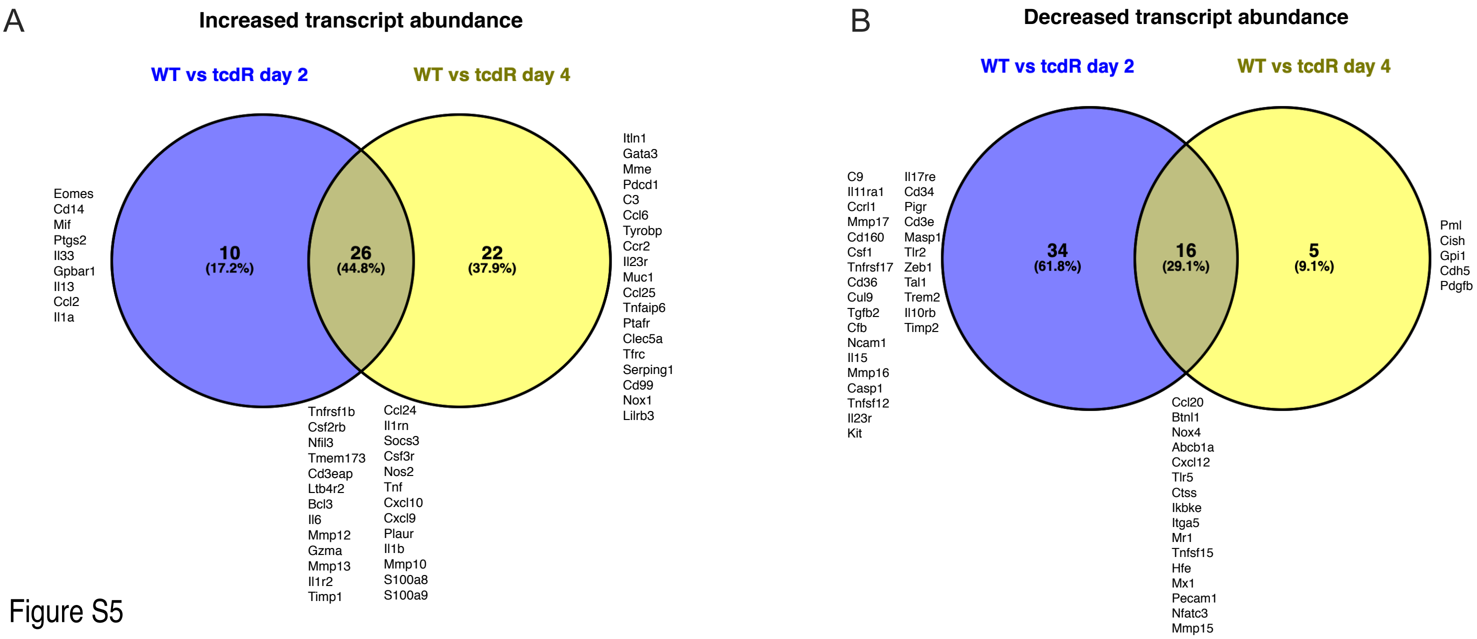

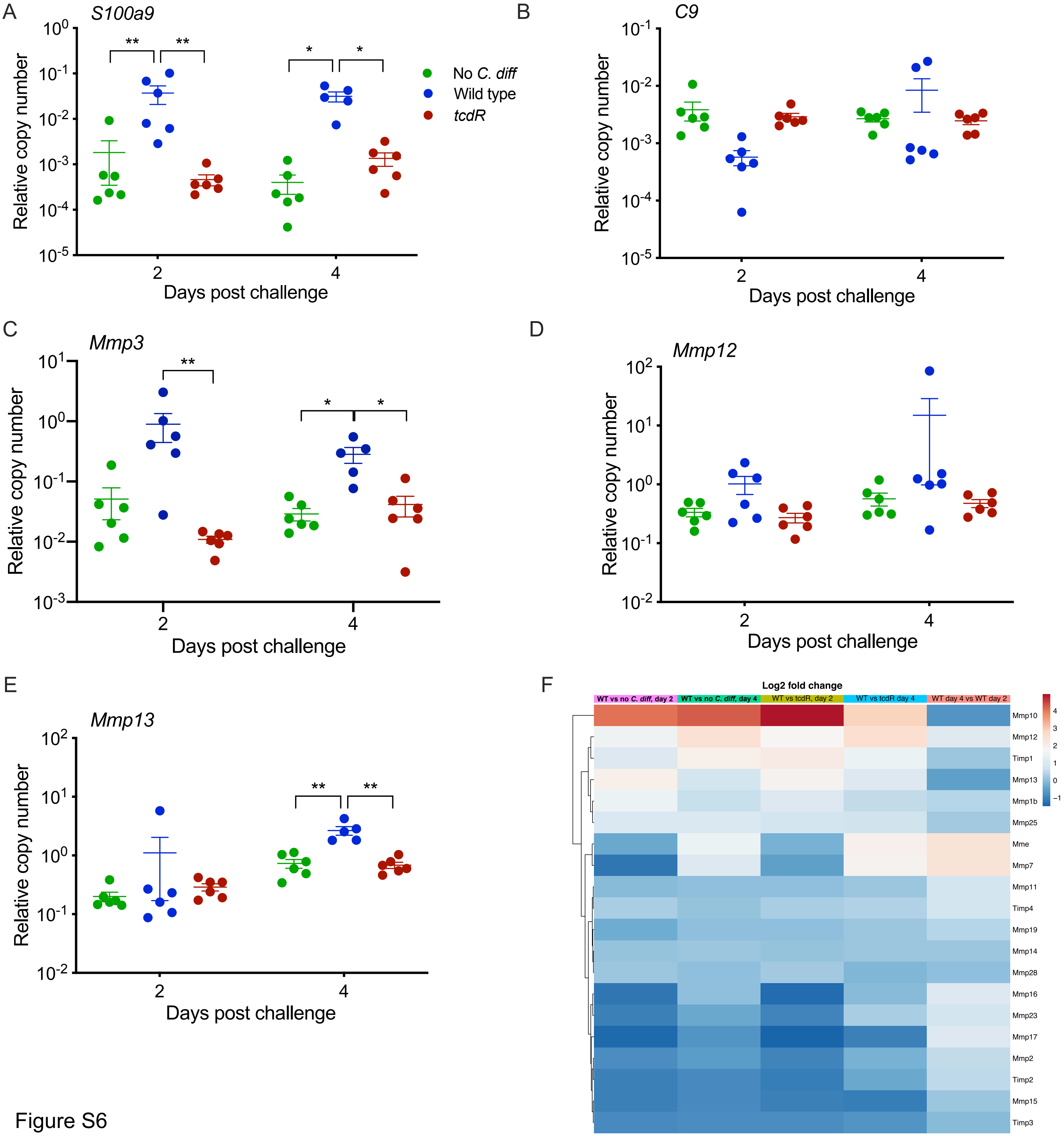
