## Supplemental File 9 for "*Clostridioides difficile* exploits toxin-mediated inflammation to alter the host nutritional landscape and exclude competitors from the gut microbiota"

### Principal components analysis from QIIME2 profiles

Michael McLaren

2019-11-01

##### Setup

```
library(here)
#> here() starts at /home/michael/ncsu-drive/research/fletcher-foley
library(phyloseq)
library(biomformat)
library(tidyverse)
#> -- Attaching packages --------------------------------------- tidyverse 1.3.0 --
#> v ggplot2 3.3.2     v purrr   0.3.4
#> v tibble  3.0.1     v dplyr   1.0.0
#> v tidyr   1.1.0     v stringr 1.4.0
#> v readr   1.3.1     v forcats 0.5.0
#> -- Conflicts ------------------------------------------ tidyverse_conflicts() --
#> x dplyr::filter() masks stats::filter()
#> x dplyr::lag()    masks stats::lag()
library(vegan)
#> Loading required package: permute
#> Loading required package: lattice
#> This is vegan 2.5-6
library(cowplot) # Only needed for cowplot theme
#>
#> ********************************************************
#> Note: As of version 1.0.0, cowplot does not change the
#>   default ggplot2 theme anymore. To recover the previous
#>   behavior, execute:
#>   theme_set(theme_cowplot())
#> ********************************************************
library(ggrepel) # Only needed for taxa labels in PCA plot
library(janitor)
#>
#> Attaching package: 'janitor'
#> The following objects are masked from 'package:stats':
#>
#>     chisq.test, fisher.test
# Also used: Biostrings, broom, knitr
```

```r
theme_set(theme_cowplot(font_size = 12))
```

```r
# otu table -----------------------------------------------------------------
# Read in the biom file without extracting the entire qza file
qza_file <- here("data", "fletcher", "qiime2", "table.qza")
# create a new directory
tmp <- tempfile()
dir.create(tmp)
# Get the path to the fasta file within the qza object
flist <- unzip(qza_file, list = TRUE) %>%
    as_tibble
biom_file <- flist %>%
    filter(str_detect(Name, "\\.biom")) %>%
    pull(Name)
# unzip and load just the fasta file
unzip(qza_file, files = biom_file, exdir = tmp)
bm <- file.path(tmp, biom_file) %>%
    read_biom()
#> Warning in strsplit(conditionMessage(e), "\n"): input string 1 is invalid in
#> this locale
# Make otu table
otu <- biom_data(bm) %>%
    as("matrix") %>%
    otu_table(taxa_are_rows = TRUE)
dim(otu)
#> [1] 208  32
# reference sequences -------------------------------------------------------
# Read in the reference sequences files without extracting the entire qiime qza
qza_file <- here("data", "fletcher", "qiime2", "rep-seqs.qza")
# create a new directory
# tmp <- dir_create(file_temp())
tmp <- tempfile()
dir.create(tmp)
# Get the path to the fasta file within the qza object
flist <- unzip(qza_file, list = TRUE) %>%
    as_tibble
fasta_file <- flist %>%
    filter(str_detect(Name, "\\.fasta")) %>%
    pull(Name)
# unzip and load just the fasta file
unzip(qza_file, files = fasta_file, exdir = tmp)
rs <- file.path(tmp, fasta_file) %>%
    Biostrings::readDNAStringSet()
# sample metadata -----------------------------------------------------------
trts.fletcher <- c("cef", "wt", "tcdR")
cages.fletcher <- seq(200, 211)
sam <- tibble(sample = sample_names(otu)) %>%
    separate(sample, c("cage", "mouse", "day", "treatment"), sep = "-",
        remove = FALSE) %>%
    mutate(day = str_extract(day, "[0-9]+")) %>%
    mutate_at(vars(cage, mouse, day), as.integer) %>%
    mutate_at(vars(cage, mouse, day, treatment), as.factor) %>%
    mutate(treatment = fct_relevel(treatment, trts.fletcher)) %>%
```

```r
    column_to_rownames("sample") %>%
    sample_data
head(sam)
#>               cage mouse day treatment
#> 200-2-d4-cef  200     2   4       cef
#> 200-3-d4-cef  200     3   4       cef
#> 201-1-d2-cef  201     1   2       cef
#> 201-2-d4-cef  201     2   4       cef
#> 201-3-d4-cef  201     3   4       cef
#> 202-1-d4-cef  202     1   4       cef
tail(sam)
#>                cage mouse day treatment
#> 209-3-d4-tcdR  209     3   4      tcdR
#> 210-1-d4-tcdR  210     1   4      tcdR
#> 210-2-d2-tcdR  210     2   2      tcdR
#> 210-3-d4-tcdR  210     3   4      tcdR
#> 211-1-d2-tcdR  211     1   2      tcdR
#> 211-3-d2-tcdR  211     3   2      tcdR
# taxonomy ---------------------------------------------------------------------
qza_file <- here("data", "fletcher", "qiime2", "taxonomy.qza")
tmp <- tempfile()
dir.create(tmp)
flist <- unzip(qza_file, list = TRUE) %>%
    as_tibble
tax_file <- flist %>%
    filter(str_detect(Name, "taxonomy.tsv")) %>%
    pull(Name)
unzip(qza_file, files = tax_file, exdir = tmp)
# load and parse the taxonomy tsv file
rns <- c("kingdom", "phylum", "class", "order", "family", "genus", "species")
tax <- file.path(tmp, tax_file) %>%
    read_tsv(comment = "#") %>%
    clean_names %>%
    separate(taxon, rns, sep = ";", fill = "right") %>%
    mutate_at(rns, str_sub, 6) %>%
    select(-confidence) %>%
    column_to_rownames("feature_id") %>%
    as("matrix") %>%
    tax_table
#> Parsed with column specification:
#> cols(
#>   `Feature ID` = col_character(),
#>   Taxon = col_character(),
#>   Confidence = col_double()
#> )
# Make phyloseq object (with taxa as columns) ---------------------------------
ps <- phyloseq(otu, sam, rs, tax)
if (taxa_are_rows(ps))
    ps <- t(ps)
remove(otu, sam, rs, tax)
# Check in decreasing order
taxa_sums(ps) %>%
    all.equal(., sort(., decreasing = TRUE))
```

```
#> [1] TRUE
# Simplify taxa names
taxa_names(ps) <- paste0("ASV", seq(ntaxa(ps)))
# save for future use
saveRDS(ps, here("results", "fletcher-qiime2-phyloseq.Rds"))
```

##### Aggregate highly similar ASVs

```
d <- refseq(ps) %>%
    Biostrings::stringDist(method = "levenshtein")
Biostrings::width(refseq(ps)) %>% summary
#>    Min. 1st Qu.  Median    Mean 3rd Qu.    Max.
#>   252.0   253.0   253.0   252.9   253.0   254.0
# tree <- hclust(d) %>% as.phylo
hc <- hclust(d, method = "complete")
# plot(hc, hang=-1)
```

Form clusters where the distance between the furthest pair is < 1%, or ~2.5, in an attempt to aggregate ASVs that are 99% similar.

```
ct <- cutree(hc, h = 253*0.01) %>% enframe("taxon", "group")
# ct <- cutree(hc, h = 3) %>% enframe("taxon", "group")
archetypes <- taxa_sums(ps) %>%
    enframe("taxon", "sum") %>%
    left_join(ct, by = "taxon") %>%
    group_by(group) %>%
    top_n(1, sum) %>%
    slice(1) %>%
    select(group, archetype = taxon)
tb <- otu_table(ps) %>% as("matrix") %>% as_tibble(rownames = "sample") %>%
    gather("taxon", "abundance", -sample) %>%
    left_join(ct, by = "taxon") %>%
    group_by(sample, group) %>%
    summarize_at("abundance", sum) %>%
    left_join(archetypes, by = "group")
newotu <- tb %>%
    select(-group) %>%
    pivot_wider(names_from = archetype, values_from = abundance) %>%
    column_to_rownames("sample") %>%
    otu_table(taxa_are_rows = FALSE)
ps0 <- ps
otu_table(ps0) <- newotu
```

Check that the distances between groups are as expected. In particular, the minimum complete-linkage distance between any two groups should be 3.

```
dist_groups <- function(d, g, diag = FALSE, upper = TRUE) {
    gtb <- tibble(item = labels(d) %>% factor(., .), group = factor(g))
    dtb <- d %>%
        broom::tidy(diag = diag, upper = upper) %>%
        left_join(gtb, by = c("item1" = "item")) %>%
        left_join(gtb, by = c("item2" = "item"), suffix = c("1", "2")) %>%
        select(-distance, distance)
}
```

```
dtb <- dist_groups(d, ct$group)
# The complete-linkage dist between groups is the max dist between all pairs of
# items from the two groups; compute and then get the min dist between groups
dtb %>%
    group_by(group1, group2) %>%
    summarize_at("distance", max) %>%
    filter(group1 != group2) %>%
    pull(distance) %>%
    min
#> [1] 3
```

Check for tax concordance among the merged groups:

```
tax <- tax_table(ps) %>% as("matrix") %>% as_tibble(rownames = "taxon")
# left_join(ct, archetypes, by = "group")
tb <- ct %>%
    left_join(archetypes, by = "group") %>%
    left_join(tax, by = "taxon") %>%
    unite(taxonomy, rns, sep = ";") %>%
    select(archetype, taxonomy) %>%
    group_by(archetype) %>%
    distinct %>%
    count
#> Note: Using an external vector in selections is ambiguous.
#> i Use `all_of(rns)` instead of `rns` to silence this message.
#> i See <https://tidyselect.r-lib.org/reference/faq-external-vector.html>.
#> This message is displayed once per session.
tb %>% pull(n) %>% summary
#>    Min. 1st Qu.  Median    Mean 3rd Qu.    Max.
#>   1.000   1.000   1.000   1.024   1.000   2.000
tb %>% filter(n > 1) %>%
    left_join(tax, by = c(archetype = "taxon"))
#> # A tibble: 4 x 9
#> # Groups:   archetype [4]
#>   archetype     n kingdom  phylum  class   order  family   genus      species
#>   <chr>     <int> <chr>    <chr>   <chr>   <chr>  <chr>    <chr>      <chr>
#> 1 ASV40         2 Bacteria Firmic~ Clostr~ Clost~ Lachnos~ Lachnospi~ <NA>
#> 2 ASV43         2 Bacteria Firmic~ Clostr~ Clost~ Lachnos~ Lachnoclo~ <NA>
#> 3 ASV66         2 Bacteria Firmic~ Clostr~ Clost~ Ruminoc~ uncultured <NA>
#> 4 ASV93         2 Bacteria Firmic~ Clostr~ Clost~ Lachnos~ Lachnospi~ unculture~
```

None of the taxa with non-concordant taxonony are labeled in the plot below, so we don't need to worry about adding NAs at their lower ranks.

#### Make the plot

```
set.seed(43)
# ps0 <- ps
sam <- sample_data(ps0) %>% as("data.frame") %>% as_tibble(rownames = "sample")
tax <- tax_table(ps) %>% as("matrix") %>% as_tibble(rownames = "taxon")
# Hellinger PCA ------------------------------------------------------------
ps0.hel.pca <- ps0 %>%
    transform_sample_counts(function(x) sqrt(x / sum(x))) %>%
    {rda(otu_table(.))}
```

```r
# Get a tibble for plotting
ps0.hel.pca.tb <- scores(ps0.hel.pca, 1:4, scaling = "sites") %>%
    # merge species and sites (samples) into a single dataframe
    map_dfr(as_tibble, rownames = "var", .id = "type") %>%
    # join with species and site metadata
    left_join(sam, by = c(var = "sample")) %>%
    left_join(tax, by = c(var = "taxon")) %>%
    mutate_at("type", fct_recode, taxon = "species", sample = "sites")
# percent variance explained, formatted for plotting
ps0.hel.pca.perc <- ps0.hel.pca %>% eigenvals %>% {. / sum(.)} %>%
    {round(100 * ., 0)}
# Data frame with taxonomy info for the labeled taxa. Pick top 10 taxa based on
# distance from the origin
tax.lab <- ps0.hel.pca.tb %>%
    filter(type == "taxon") %>%
    mutate(distance = sqrt(PC1^2 + PC2^2)) %>%
    top_n(10, distance) %>%
    select(taxon = var) %>%
    left_join(tax, by = "taxon") %>%
    arrange(str_sub(taxon, 4) %>% as.integer)
ps0.hel.pca.plt <- ps0.hel.pca.tb %>%
    mutate(PC2 = -PC2) %>%
    ggplot(aes(PC1, PC2)) +
    # axes
    geom_hline(yintercept = 0, color = "black") +
    geom_vline(xintercept = 0, color = "black") +
    # all taxa
    geom_point(data = ~filter(., type == "taxon"),
        aes(PC1/3, PC2/3),
        shape = 3, color = "darkgrey") +
    # samples
    geom_point(data = ~filter(., type == "sample"),
        aes(color = treatment, shape = day), size = 3) +
    # label subset of taxa
    geom_text_repel(data = ~filter(., var %in% tax.lab$taxon),
        aes(PC1/3, PC2/3, label = var), color = "black") +
    # geom_text_repel(data = ~filter(., type == "sample"), aes(label = cage)) +
    scale_color_brewer(type = "qual", palette = 2) +
    theme(axis.line = element_blank()) +
    labs(
        title = "PCA on Hellinger-transformed abundances",
        x = paste0("PC1 [", ps0.hel.pca.perc["PC1"], "%]"),
        y = paste0("PC2 [", ps0.hel.pca.perc["PC2"], "%]")
    )
```

```r
ps0.hel.pca.plt
```

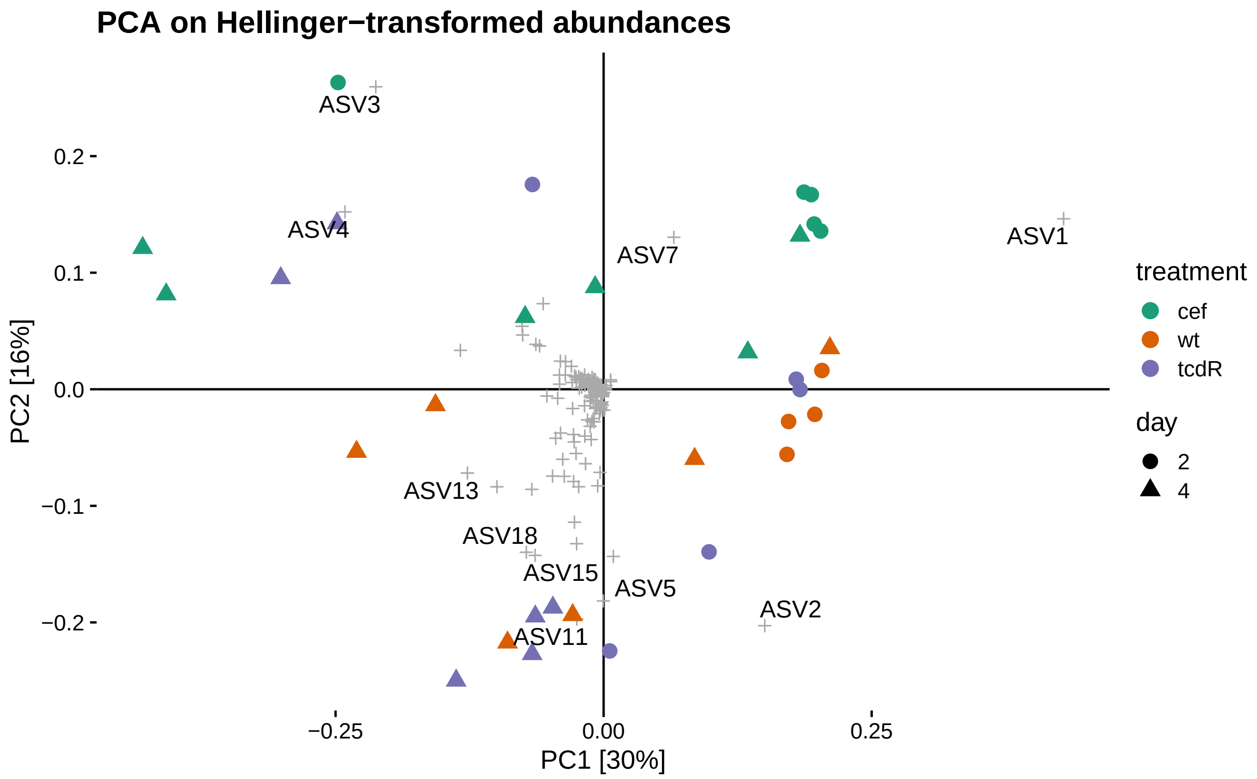

```r
ggsave(here("figures", "2019-11-5-qiime-microbiota-hellinger-pca.pdf"),
       useDingbats = FALSE, units = "in", width = 8, height = 5)
```

```r
tax.lab %>% knitr::kable()
```

| taxon | kingdom | phylum | class | order | family | genus | species |
|---|---|---|---|---|---|---|---|
| ASV1 | Bacteria | Firmicutes | Bacilli | Lactobacillales | Enterococcaceae | Enterococcus | NA |
| ASV2 | Bacteria | Firmicutes | Clostridia | Clostridiales | Peptostreptococcaceae | Clostridioides | NA |
| ASV3 | Bacteria | Bacteroidetes | Bacteroidia | Bacteroidales | Bacteroidaceae | Bacteroides | NA |
| ASV4 | Bacteria | Verrucomicrobia | Verrucomicrobiae | Verrucomicrobiales | Akkermansiaceae | Akkermansia | NA |
| ASV5 | Bacteria | Firmicutes | Clostridia | Clostridiales | Lachnospiraceae | ASF356 | NA |
| ASV7 | Bacteria | Firmicutes | Bacilli | Bacillales | Staphylococcaceae | Staphylococcus | NA |
| ASV11 | Bacteria | Firmicutes | Clostridia | Clostridiales | Lachnospiraceae | NA | NA |
| ASV13 | Bacteria | Firmicutes | Clostridia | Clostridiales | Lachnospiraceae | NA | NA |
| ASV15 | Bacteria | Firmicutes | Erysipelotrichia | Erysipelotrichales | Erysipelotrichaceae | uncultured | uncultured bacterium |
| ASV18 | Bacteria | Firmicutes | Clostridia | Clostridiales | Lachnospiraceae | NA | NA |

```r
dna <- refseq(ps) %>% as.character %>% enframe("taxon", "sequence")
tax.lab %>%
    left_join(dna, by = "taxon") %>%
    write_csv(here("results", "2019-11-5-qiime-ordination-taxa.csv"))
```

#### Session info

```r
sessionInfo()
#> R version 4.0.2 (2020-06-22)
```

```
#> Platform: x86_64-pc-linux-gnu (64-bit)
#> Running under: Arch Linux
#>
#> Matrix products: default
#> BLAS:   /usr/lib/libblas.so.3.9.0
#> LAPACK: /usr/lib/liblapack.so.3.9.0
#>
#> locale:
#>  [1] LC_CTYPE=en_US.UTF-8       LC_NUMERIC=C
#>  [3] LC_TIME=en_US.UTF-8        LC_COLLATE=en_US.UTF-8
#>  [5] LC_MONETARY=en_US.UTF-8    LC_MESSAGES=en_US.UTF-8
#>  [7] LC_PAPER=en_US.UTF-8       LC_NAME=C
#>  [9] LC_ADDRESS=C               LC_TELEPHONE=C
#> [11] LC_MEASUREMENT=en_US.UTF-8 LC_IDENTIFICATION=C
#>
#> attached base packages:
#> [1] stats     graphics  grDevices utils     datasets  methods   base
#>
#> other attached packages:
#>  [1] janitor_2.0.1     ggrepel_0.8.2     cowplot_1.0.0     vegan_2.5-6
#>  [5] lattice_0.20-41   permute_0.9-5     forcats_0.5.0     stringr_1.4.0
#>  [9] dplyr_1.0.0       purrr_0.3.4       readr_1.3.1       tidyr_1.1.0
#> [13] tibble_3.0.1      ggplot2_3.3.2     tidyverse_1.3.0   biomformat_1.16.0
#> [17] phyloseq_1.32.0   here_0.1          rmarkdown_2.3     nvimcom_0.9-92
#> [21] usethis_1.6.1
#>
#> loaded via a namespace (and not attached):
#>  [1] nlme_3.1-148        fs_1.4.1            lubridate_1.7.9
#>  [4] RColorBrewer_1.1-2  httr_1.4.1          rprojroot_1.3-2
#>  [7] tools_4.0.2         backports_1.1.6     utf8_1.1.4
#> [10] R6_2.4.1            DBI_1.1.0           BiocGenerics_0.34.0
#> [13] mgcv_1.8-31         colorspace_1.4-1    ade4_1.7-15
#> [16] withr_2.2.0         tidyselect_1.1.0    compiler_4.0.2
#> [19] cli_2.0.2           rvest_0.3.5         Biobase_2.48.0
#> [22] xml2_1.3.2          labeling_0.3        scales_1.1.1
#> [25] digest_0.6.25       XVector_0.28.0      pkgconfig_2.0.3
#> [28] htmltools_0.5.0     highr_0.8           dbplyr_1.4.4
#> [31] rlang_0.4.6         readxl_1.3.1        rstudioapi_0.11
#> [34] farver_2.0.3        generics_0.0.2      jsonlite_1.7.0
#> [37] magrittr_1.5        Matrix_1.2-18       fansi_0.4.1
#> [40] Rcpp_1.0.4.6        munsell_0.5.0       S4Vectors_0.26.1
#> [43] Rhdf5lib_1.10.0     ape_5.4             lifecycle_0.2.0
#> [46] stringi_1.4.6       yaml_2.2.1          snakecase_0.11.0
#> [49] MASS_7.3-51.6       zlibbioc_1.34.0     rhdf5_2.32.1
#> [52] plyr_1.8.6          grid_4.0.2          blob_1.2.1
#> [55] parallel_4.0.2      crayon_1.3.4        Biostrings_2.56.0
#> [58] haven_2.3.1         splines_4.0.2       multtest_2.44.0
#> [61] hms_0.5.3           knitr_1.29          pillar_1.4.4
#> [64] igraph_1.2.5        reshape2_1.4.4      codetools_0.2-16
#> [67] stats4_4.0.2        reprex_0.3.0        glue_1.4.1
#> [70] evaluate_0.14       data.table_1.12.8   modelr_0.1.8
#> [73] vctrs_0.3.1         foreach_1.5.0       cellranger_1.1.0
#> [76] gtable_0.3.0        assertthat_0.2.1    xfun_0.15
```

```
#> [79] broom_0.5.6         survival_3.1-12     iterators_1.0.12
#> [82] IRanges_2.22.2      cluster_2.1.0       ellipsis_0.3.1
```
