## Supplementary material for "*Clostridioides difficile* exploits toxin-mediated inflammation to alter the host nutritional landscape and exclude competitors from the gut microbiota": Table 1

| Primers 5’ to 3’ (lowercase for restriction sites) | Target |
| --- | --- |
| 5’tcdR.up.bamHI – ggatccTATATGAAAGAAGAGCATAATTTACCAG  3’tcdR.up.SOE - TCATTAATTACATAAAATCATCCTCTCTTATATTTATAATG | *tcdR* upstream homology arm |
| 5’tcdR.down.SOE -GGATGATTTTATGTAATTAATGAATTTAAAGAAATATTTACAATAG  3’tcdR.down.kpnI – ggtaccATATACACCACCAACTTCTTTTAAGGC | *tcdR* downstream homology arm |
| 5’tcdRc.bamh1 – ggatccGATTTCATAAAAGATACTATTTTAGTCTTG  3’tcdRc.kpn1 – ggtaccGTTAATTCTAAAATTTGATTTCTATTG | Check for loss of *tcdR* coding sequence |
| tcdR.left.up.gDNA – CAATGTTAGAAAATCATTTGAGTG  YN4.kpn.side – ATGACCATGATTACGAATTCG | Check for knockout plasmid integration |
| YN4.bamHI.side – GCGTGACGTCGACTCTAG  tcdR.right.down.gDNA - ATATCAAAATGCTCTGAAGTATATCC | Check for knockout plasmid integration |
| 5’mB.actin.intron1 - GCCTTCTTTTGTGTCTTGATAG  3’mB.actin.exon3 – CTGGGTCATCTTTTCACG | Check for host gDNA |
| 5’plaur.q – GGCACAGCAGGTTTCCATA  3’plaur.q CGGTGGAAAGCTCTGAAGAT | *Plaur* |
| 5’S100a9.q – CTCCTCAAAGCTCAGCTGATTG  3’S100a9.q – AATGGTGGAAGCACAGTTGG | *S100a9* |
| 5’c9.q – CCTGAAAGAGAAGATTCTCAGAGG  3’c9.q – CGTTTGCCAGGGACGAG | *C9* |
| 5’tlr5.q – CGCCTCATCTCACTGCATAC  3’tlr5.q - ACAGATGTGTCTGGCATATGTT | *Tlr5* |
| 5’mmp3.q – GGGATGATGATGCTGGTATG  3’mmp3.q – TGACAATAAGACTACTGTCCTTT | *Mmp3* |
| 5’mmp12.q – CTAGAAGCAACTGGGCAACT  3’mmp12.q – GCTCTAAGATGCTGTACATCGG | *Mmp12* |
| 5’mmp13.q – CTACCCACTTGTTCTAATGACCTAT  3’mmp13.q - GCTGTGTCTTAGCTGGATCTAC | *Mmp13* |
| 5’rpoC.qPCR – TGGCAGTCCATGTACCTTTATC  3’rpoC.qPCR - GGTGAACCATCTTTAGGAGCA | *rpoC* |
| 5’cdd3.q – ATCCAGATATGTTAGGTGGATTTGA  3’cdd3.q – AGCTTCCATGTATCTCCTTTATGT | *cdd3* |
| 5’PTS_fms.q – AGAATTGTCCAGAGAGCTTGTT  3’PTS_fms.q – TCTCAGCATCATCTGCTGATTTA | *PTS IIA* |
| 5’ilvD.q – CATGAACTCTTGTCCTGGATGT  3’ilvD.q – GTGACGCAGCAGTACCATTA | *ilvD* |
| 5’cat1.q TGGATTTACACCTTCAGGCTATC  3’cat1.q - CTCCATCAACTTCTGGACCTAAA | *Cat1* |
| 5’ CD630DERM_34420.q – GCCACATGGAAGACCTACAA  3’ CD630DERM_34420.q – ACGAGTCATGTCTGATTGATACC | *CD630DERM_34420* |
| 5’tcdA.qPCR – ACTAGACGAACATGACCCATTAC  3’tcdA.qPCR - GCTACCGTTGCAGCTATAGATAA | *tcdA* |
